# Discovery of tumoricidal DNA oligonucleotides by effect-directed *in-vitro* evolution

**DOI:** 10.1101/629105

**Authors:** Noam Mamet, Yaniv Amir, Erez Lavi, Liron Bassali, Gil Harari, Itai Rusinek, Nir Skalka, Elinor Debby, Mor Greenberg, Adva Zamir, Anastasia Paz, Neria Reiss, Gil Loewenthal, Irit Avivi, Avichai Shimoni, Guy Neev, Almogit Abu-Horowitz, Ido Bachelet

**Affiliations:** Augmanity, Rehovot, Israel Website: http://augm.com; Faculty of Life Sciences, Bar-Ilan University, Ramat-Gan, Israel; Aummune, Tel Aviv Sourasky Medical Center, Tel Aviv, Israel Website: http://aummune.com; Tel Aviv Sourasky Medical Center, Sackler Faculty of Medicine, Tel Aviv University, Tel Aviv, Israel; BMT Department, Division of Hematology, Sheba Medical Center Tel Hashomer, Ramat-Gan, Israel

## Abstract

Our current model of drug discovery is challenged by the relative ineffectiveness of drugs against highly variable and rapidly evolving diseases and their relatively high incidence of adverse effects due to poor selectivity. Here we describe a robust and reproducible platform which could potentially address these limitations. The platform enables rapid, *de-novo* discovery of DNA oligonucleotides evolved *in-vitro* to exert specific biological effects on target cells. Unlike aptamers, which are selected by their ligand binding capacity, this platform is driven directly by therapeutic effect and selectivity towards target vs negative target cells. The process could, therefore, operate without any *a-priori* knowledge (e.g. mutations, biomarker expression, or known drug resistance) of the target. We report the discovery of DNA oligonucleotides with direct and selective cytotoxicity towards several tumor cell lines as well as primary, patient-derived solid and hematological tumors, some with chemotherapy resistance. Oligonucleotides discovered by this platform exhibited favorable biodistribution in animals, persistence in target tumors up to 48 hours after injection, and safety in human blood. These oligonucleotides showed remarkable efficacy *in-vivo* as well as *ex-vivo* in freshly obtained, 3D cultured human tumors resistant to multiple chemotherapies. With further improvement, these findings could lead to a drug discovery model which is target-tailored, mechanism-flexible, and nearly on-demand.

## Introduction

Effect and selectivity are essential requirements for therapeutic molecules. However, it has become increasingly clear that for many severe diseases, achieving these requirements could be challenging. The continual emergence of drug resistance in cancer, for example, makes therapeutic targeting extremely difficult^1-2,3^. The problem is compounded by the high variability and patient heterogeneity of the disease^4^, making it challenging for a single drug or protocol to be both effective and safe across many patients^5^. New drugs continue to be developed despite known resistance to them and the prediction that they will be effective only for a small fraction of patients^5,6^. The current premise of personalized medicine typically refers to predicting or validating responses to drugs from the set of currently available ones^7^, leaving the problems of emergent resistance and off-target toxicity of these drugs unaddressed. Although superior to older generation chemotherapy in many ways, antibodies are specific to their antigens and would show selectivity only when antigen expression is limited to a specific target cell. Recently approved chimeric antigen receptor (CAR)-T cell therapies, while new and promising, have often shown adverse effects due to this fact^8^.

For the purposes of this study we use cancer as a case study, and argue that an effective and viable therapeutic strategy for this disease would have to satisfy three requirements:

1. It needs to be tailored to a specific tumor/patient, due to the observed variability between individual cases;
2. It needs to be selective, to minimize adverse effects or eliminate them completely; and
3. Its discovery needs to be rapid and economically repeatable, to counter the emergence of resistance.

In this article, we describe a platform that essentially fulfills these requirements. While further development and improvement are necessary to expand it and establish its clinical potential in cancer and other conditions, we report extremely promising results that should motivate this effort. This platform is based on the *in-vitro* evolution of oligonucleotides driven directly by a therapeutic effect.

The ability to artificially evolve and select nucleic acid molecules with specific properties has been known for nearly three decades^9,10^ and has produced diverse functions^11–15^, molecular and cellular specificities^16–18^, and therapeutic effects^19–21^. The SELEX^15,16^ (systematic evolution of ligands by exponential enrichment) method is routinely used to find aptamers - RNA or DNA oligonucleotides with the ability to bind a specific molecular or cellular target. In SELEX, iterating rounds of selection are applied to an initial population of 10^9^-10^15^ oligonucleotides. Selection pressure drives this population towards a subpopulation enriched with oligonucleotides capable of binding the target presented to them. The process is designed such that the oligonucleotides that are best binders survive each round and are passed on to the next one. Stringent washing steps are applied as a selective force for the removal of oligos that do not bind the target. At the end of the process, binding candidates are selected and tested separately. Importantly, previously described cell-specific aptamers have been reported to also have a secondary function subsequent to the binding. This has been usually achieved by an additional step in which the positive binders are tested again, this time at the cell level, for the secondary function^22–24^. This is done in a low throughput manner, separately testing each candidate.

The platform we describe here aims at achieving this goal and selects oligonucleotides with defined therapeutic functions, such as target cell apoptosis. The platform screens the oligonucleotide pool for the desired function, looking only for a chosen biological effect at the live cell level. A remarkable consequence of this feature is that the platform does not require any *a-priori* biological knowledge about the target, such as which cell type it is, which surface markers it expresses, which mutations its genome or epigenome carry, and which current chemotherapies it is already resistant to. The platform only requires a clear identification of the input cells as the target or as a negative target.

## Results

The platform’s workflow consists of two stages. The purpose of the first stage is to enrich a random initial single-stranded oligonucleotide library (∼10^15^) for specific target binders, however without significant reduction of its diversity. This is done by 3 rounds of a conventional cell-SELEX process **(Supplementary note 1)**. The enriched population then exits into the second, functional stage. The crucial challenge of this stage is that candidate functional oligonucleotides distribute evenly across the target cell sample, generating a very low effective concentration and are therefore highly unlikely to generate any noticeable effect on cells. To overcome this issue and enhance the signal, we created an emulsion in which the library was dispersed together with microparticles, and performed emulsion PCR (ePCR) to coat each microparticle with multiple copies of a single oligonucleotide, or at most very few different ones. Thus each cluster creates a local high concentration for the oligonucleotide it holds **(Supplementary note 2)**. The process then starts and continues with the population of clustered oligonucleotides rather than a solution-phase library.

Each functional round commences by incubating the clustered library with target cells which were loaded with a reporter indicating a desired effect, such as apoptosis or cell activation. Following incubation, the cell:cluster mixture is sorted by FACS to isolate cluster^+^/effect^+^ events. Clusters are then eluted from cells, amplified by PCR, and amplified again by ePCR to generate new clusters for the next round. Two final analyses are then performed: first, all output libraries from the functional stage are experimentally compared in their ability to induce an effect in the target cells; this test validates that the process has successfully driven the library towards improvement. Second, analysis by deep sequencing highlights the most successful oligonucleotides for further synthesis and functional validation.

This workflow was verified on a colorectal carcinoma cell line, HCT-116. HCT-116 cells first went through 7 rounds of cell-SELEX, followed by a binding assay **(Fig. 1A, B)**. The output library from round 3 was then introduced into the functional stage for an additional 6 to 8 rounds. The functional stage is based on choosing a specific marker or mechanism for targeting by the library. As a functional reporter in this case study, we chose a fluorogenic substrate of activated caspase-3/7 (cas-3/7). This reporter produced a good signal for sorting. Sorting of cluster^+^/cas-3/7^+^ events went on for 8 rounds **(Fig. 1C)**. In each round, the incubation time for generating effect was 1.5 h. Importantly, cells entering the round being already dead are gated out based on their physical parameters, to prevent enrichment of dead cell-binding oligos, which are a potentially significant contaminant. Strikingly, a comparison of the output libraries from all functional rounds demonstrated a consistent improvement in the library’s ability to induce cas-3/7 activity in HCT-116 cells **(Fig. 1D)**.

**Figure 1.**
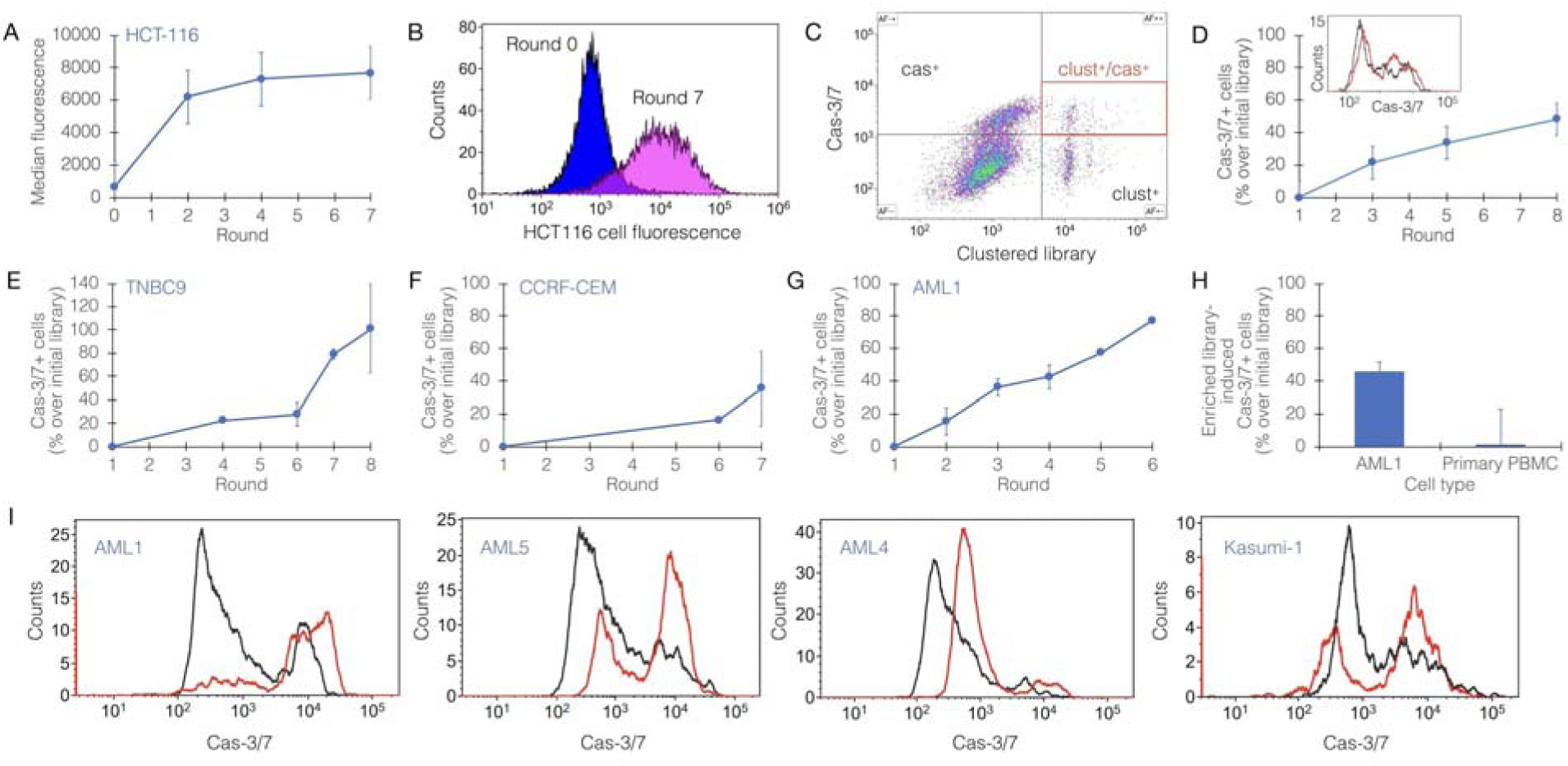
Tumoricidal oligonucleotide libraries created by effect-directed *in-vitro* evolution. **A**, consistent enrichment of the binding capacity of a random library during the initial stage of the process, implemented on HCT116, a colorectal cancer cell line. Each point represents an independent binding assay. **B**, a representative binding assay showing the success of the initial stage (round 7 [final] vs round 0 of the process) on HCT116 cells. **C**, representative sorting plot during the functional stage of the process. Bead-clustered library (x-axis) is tagged by Cy5; Cas-3/7 is a green fluorescent reporter. Red gate within the upper right quadrant includes cas-3/7^+^ (cas^+^) cells bound to an oligo cluster (clust^+^). These events are sorted and carried forward to the next round. **D**, consistent enrichment of the ability to induce cas-3/7 activation in HCT116 cells by the oligo library. Inset shows flow cytometric analysis of cas-3/7 activity in HCT116 cells (black, cells treated with round 1; red, cells treated with round 8). **E-G**, representative runs of effect-directed *in-vitro* evolution, resulting in oligo libraries with cas-3/7 activity-inducing capacity: **E**, patient-derived xenograft (PDX)-derived triple negative breast cancer (TNBC) cells, termed TNBC9; **F**, human acute lymphoblastic leukemia cell line (CCRF-CEM); **G**, patient-derived acute myeloblastic leukemia (AML), termed AML1. **H**, the selectivity of tumoricidal oligo library towards AML1 compared with primary peripheral blood mononuclear cells (PBMC) from a healthy donor. Shown is a representative analysis of cas-3/7 activity in both targets induced by the same library. The effect observed in primary PBMC is statistically zero. **I**, the exclusivity of a library evolved against AML1 target cells, to AML cells from other patients (AML4, AML5) and an AML cell line, kasumi-1. Shown is a flow cytometric analysis of cas-3/7 activity (black, round 1; red, round 6 [final] of the process).

We repeated this workflow on several tumor targets of human origin: primary triple negative breast cancer (TNBC), an acute lymphoblastic leukemia (ALL) cell line, and primary acute myeloid leukemia (AML). TNBC cells (termed TNBC9) were produced from patient-derived xenografts as previously described^25^. MCF10A cells, a non-tumorigenic breast epithelial cell line^26^, were used as negative target cells. These runs resulted in oligonucleotide libraries which exerted potent and selective cytotoxicity on the target cells, including those derived directly from patients **(Fig. 1E, F, G, H)**. AML cells (termed AML1) were freshly isolated from patient’s blood **(Supplementary note 3)**, and peripheral blood mononuclear cells (PBMC) from a healthy donor were used as negative target cells. Here, too, the process successfully produced a library which induced cas-3/7 activation in AML cells but not in PBMC from a healthy donor **(Fig. 1G, H)**. To evaluate the exclusivity of libraries to the target cells used in their evolution, we examined the ability of the AML library described above (AML1) to induce apoptosis in other AML cells, both freshly-isolated from patients, and a cell line. Interestingly, some target cells exhibited partial resistance to this library, while others were significantly susceptible **(Fig. 1I)**.

In order to evaluate the applicability of this platform as a therapeutic strategy for cancer, we followed the TNBC9 targeting results with selection of lead molecules. Based on sequencing analysis **(Fig. 2A)**, 10 candidate oligonucleotides were selected, synthesized, and folded. The effectiveness of these candidates in target killing was measured on the PDX-derived TNBC9 cells, highlighting a single oligonucleotide, termed E8, as the most effective **(Fig. 2B)**. The observed level of direct target killing by E8 *in-vitro* and *ex-vivo* ranged between ∼ 20-40% in independent biological replicate experiments, which is comparable to the levels observed with approved anti-cancer biologicals^27–29^. E8 demonstrated remarkable selectivity at the target cell level, killing TNBC9 but not MCF10A cells, which were used as negative targets in the *in-vitro* evolution process **(Fig. 2C)**. E8 was not exclusive to TNBC9 and showed a remarkable effect on MDA-MB-231 cells as well **(Fig. 2D)**. In preparation for *in-vivo* testing, these effects were re-validated using E8 modified with poly-ethylene glycol (PEG), a modification that extends *in-vivo* stability and half-life of the oligonucleotide, demonstrating that the effect was retained with PEG **(Fig. 2E)**. In addition, E8 retained function in mouse serum **(Fig. 2F)**.

**Figure 2.**
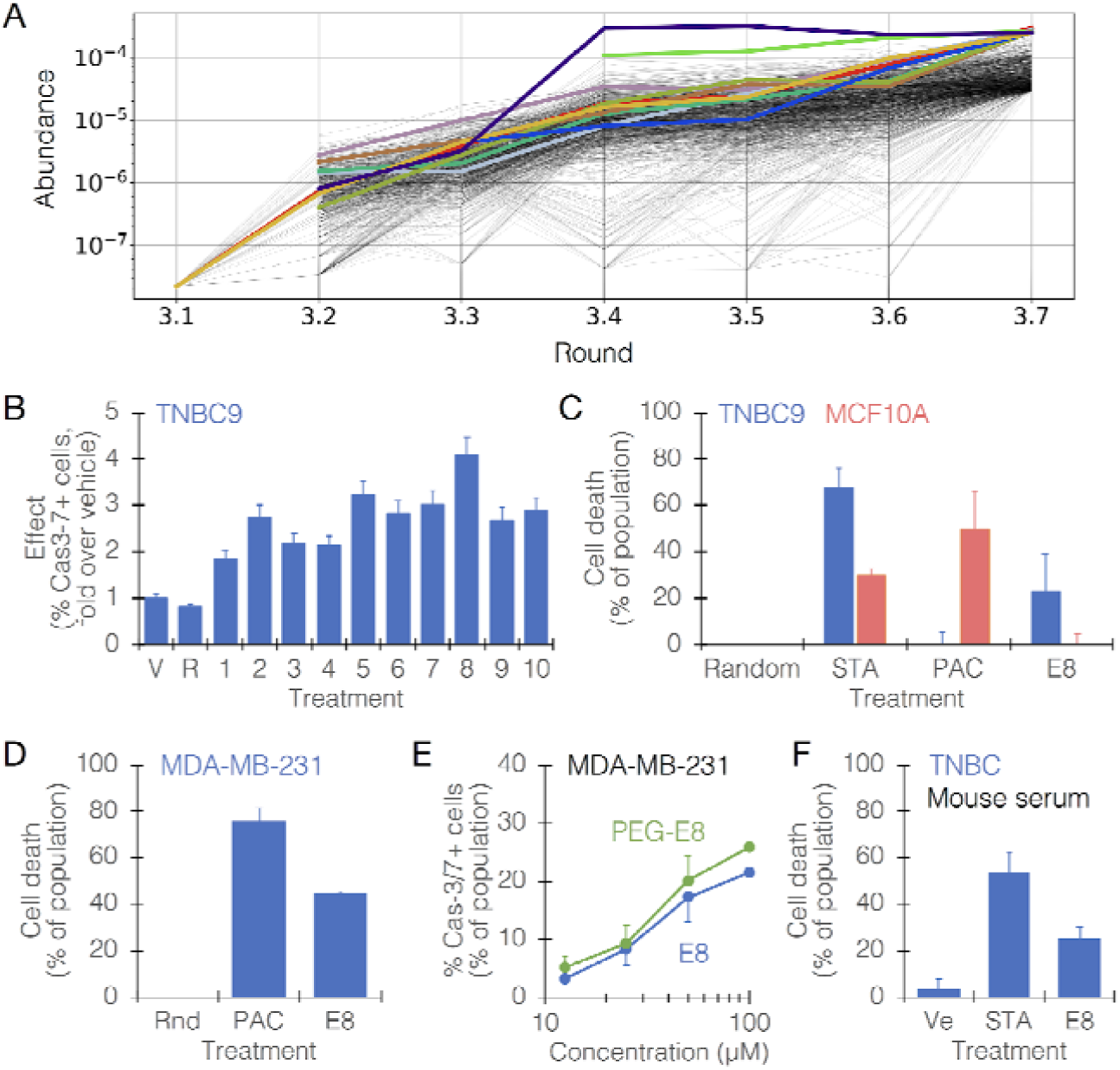
Identification of a lead candidate, E8, from a tumoricidal oligo library. Cell type are denoted in the top left corner of plots. **A**, sequence abundance plot from a representative effect-directed *in-vitro* evolution run. The plot shows a random sample of 1,000 sequences out of the 10,000 most abundant sequences in each round (traces are shown from their first appearance in the data) with the 10 most abundant ones highlighted in color. These were synthesized and screened to find candidates. **B**, a representative screening to highlight effective oligos. The response was measured as the ability to induce significant cas-3/7 activation in the population compared with vehicle. V, vehicle; R, random oligonucleotide; 1-10, oligo IDs (E1, E2,.. E10). **C**, Selectivity of E8 to TNBC9 cells (blue) over the negative target cells, MCF10A (red). The effect was measured by a cell viability count assay. STA, staurosporine; PAC, paclitaxel; Random, random oligonucleotide. **D**, the effect of E8 on MDA-MB-231 cells. Rnd, random oligonucleotide; PAC, paclitaxel. **E**, the dose-response curve for E8 (blue) and PEGylated (PEG)-E8 (green), showing persistence of effect in the modified oligo. The two curves are statistically indifferent. **F**, the effect of E8 on TNBC cells in mouse serum. Ve, vehicle; STA, staurosporine.

To determine the dispersion of E8 we used fluorescently-labeled E8 as previously described for aptamer in-vivo imaging probes^30–32^. The molecule, modified with 5’ Cy5.5 and 3’ PEG, was injected intravenously in two doses (6 and 60 mg/kg) into NOD/SCID mice in which MDA-MB-231 tumors were induced. These experiments showed that E8 localizes to and is significantly retained in the tumors at 24 and 48 h post injection **(Fig. 3A, B, C)**. Furthermore, when E8 was mixed with whole human blood from healthy donors, no hemolysis, agglutination, or cytokine responses were observed **(Supplementary note 5)**. To evaluate the efficacy of E8, the PEGylated oligonucleotide was injected once/2 days during the course of an 11-day period, at a dose of 100 mg/kg (equivalent in molar terms to standard chemotherapy). During this period, in E8-treated animals, tumor growth was inhibited, with mean tumor volumes significantly lower than in vehicle-treated animals (final volumes: 168±39 vs 301±51 mm^3^ in E8-treated animals and vehicle-treated ones, respectively) **(Fig. 3D)**. Remarkably, tumors extracted from E8-treated animals exhibited macroscopic signs of tissue death **(Fig. 3E)**. Analysis of caspase-3 activity in histological sections from these tumors showed significant staining in tumors from E8-treated animals **(Fig. 3F, G)**, reinforcing the hypothesis that this effect was caused directly by E8, which was selected from a library evolved specifically to activate caspase 3. Tissue sections were also analyzed by TUNEL showing marked effect in E8-treated tumors. Importantly, no significant changes in appearance or body weight were observed following injections **(Supplementary note 4)**.

**Figure 3.**
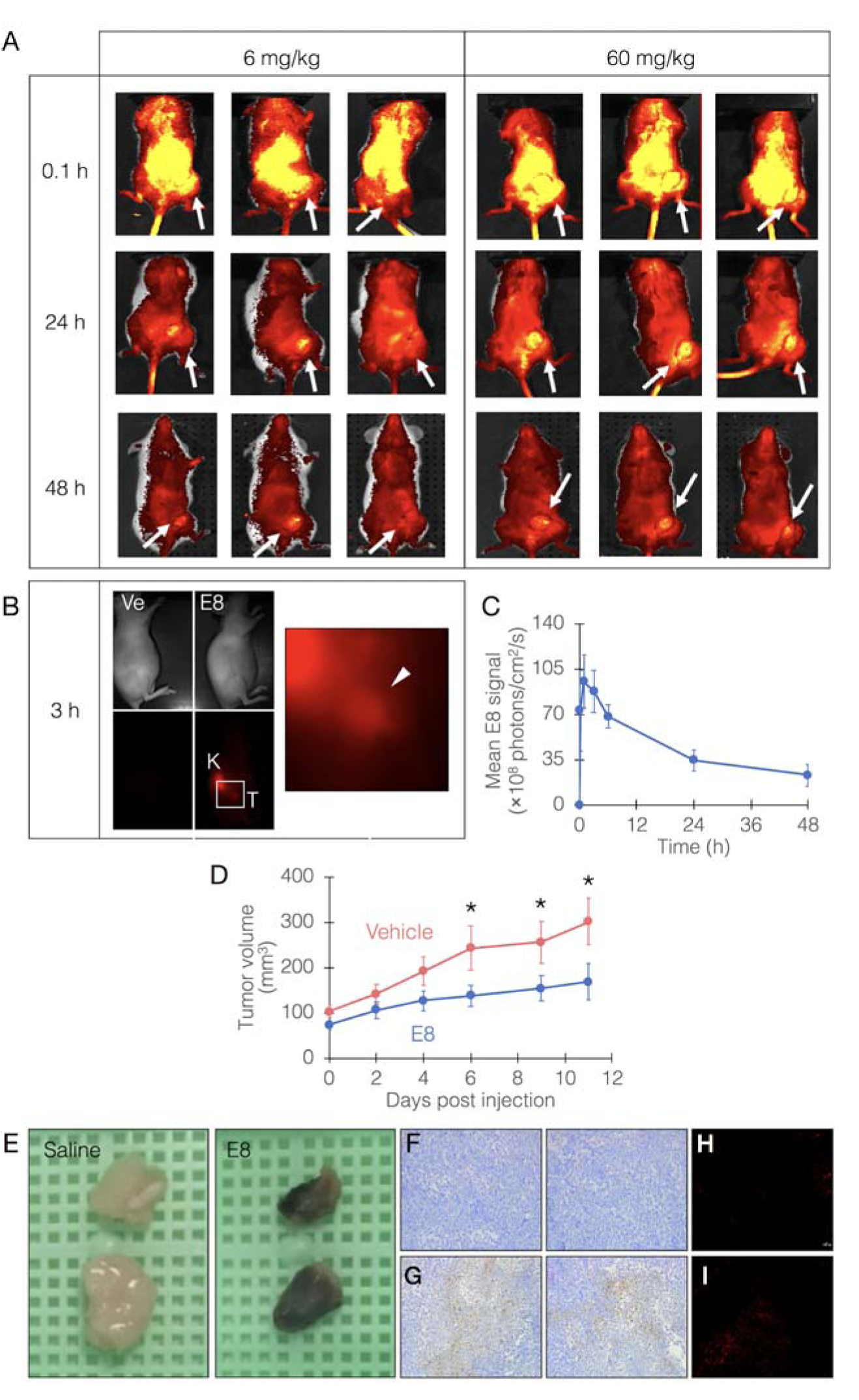
E8 biodistribution and efficacy in an animal model. **A**, E8, modified with Cy5.5 and PEG, was injected at 6 or 60 mg/kg, i.v. into NOD/SCID mice bearing MDA-MB-231-derived tumors on their right hind limb. Fluorescence was measured *in-vivo* immediately after injection and following 24 and 48 h. White arrows point to tumor locations. **B**, homing and retention of E8 at tumor site 3 h following i.v. injection (Ve, vehicle; K, kidney). Inset region is shown magnified on the right. White arrowhead points to tumor site. **C**, quantitative measurement of E8 level in tumors up to 48 h post injection. E8 level peaks at 1-3 h post injection, then fall but is still maintained later. **D**, the efficacy of E8 in mice bearing MDA-MB-231-derived tumors. E8 was injected at 100 mg/kg, 1 dose/2 d for 11 d, and tumor volumes were measured. Mean tumor volumes of the two groups are statistically indifferent at day 0. Asterisks denote a statistically significant difference with *p<*0.05 (*n* = 8 mice/group). **E**, representative photographs of tumors excised from mice sacrificed at day 11. Tumors from E8-treated mice appear necrotic. **F-G**, histochemical analysis of caspase 3 activity in tumor-derived tissue sections (F, vehicle-treated; G, E8-treated). Caspase 3 activity is exhibited as brown color. **H-I**, TUNEL analysis of tumor-derived tissue sections (H, vehicle-treated; I, E8-treated).

The efficacy of E8 was also evaluated in human *ex-vivo* organ cultures (EVOC)^7,33^ freshly derived from BC patients **(Supplementary note 6)**. The pathological assessment showed that E8 had a significant effect (grades 3-4 on a 0-4 scale) on tumor cells in the EVOC samples from 2 patients, both of them showing resistance to at least one chemotherapy **(Fig. 4A, B)**.

**Figure 4.**
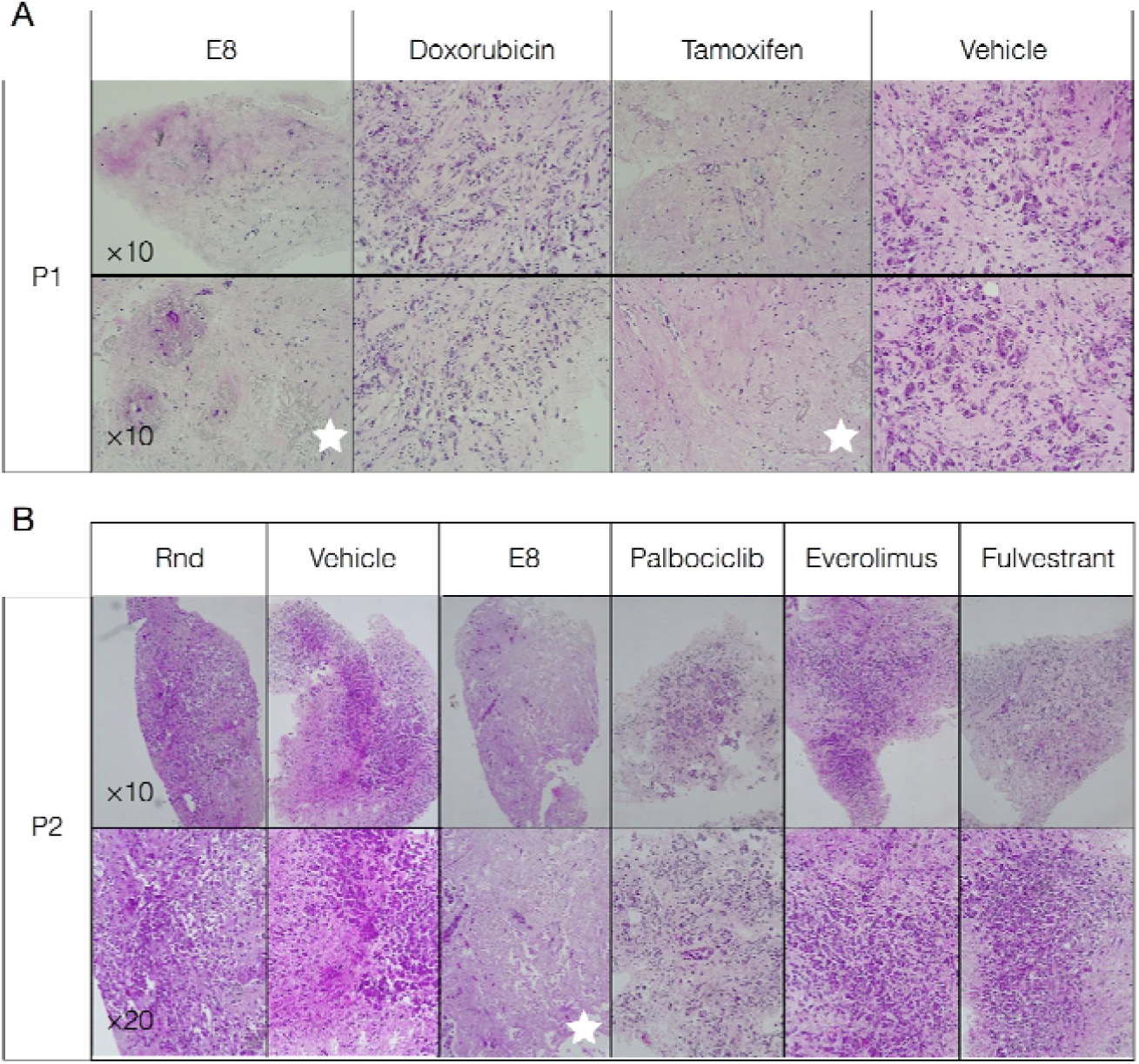
Efficacy of E8 in human *ex-vivo* organ cultures (EVOC). E8 was administered to EVOCs freshly derived from TNBC patients (two representative patients, P1 and P2, are shown). **A-B**, E8 and other chemotherapies were administered at concentrations of 20-50 LJM, 1/d for 2 d, samples were fixed at 5 d, sections were made and stained with hematoxylin-eosin. Effect were graded by 2 blinded pathologists on a 0-4 scale. White stars denote the experimental groups in which effect reached a grade of at least 3, and relate to the entire respective column. Rnd, random oligonucleotide; magnifications are shown in the bottom left corners of each row.

The described platform was reproducibly tested in *n*=9 independent runs on human tumor targets from different types and sources, with each library tested in multiple biological repeats. It is interesting to note that the platform successfully produced effective libraries against targets with known resistance to multiple drugs, suggesting that the process is driven sufficiently robustly so as to find solutions to targets following significant biological alterations (e.g. shutting down pathways to resist a drug).

## Discussion

We describe a platform for the rapid *de-novo* discovery of therapeutic oligonucleotides by effect-driven *in-vitro* evolution. This platform could potentially address the central limitations of our current model of drug discovery. Particularly, this platform receives a human sample and operates a specific algorithm to generate a new therapeutic molecule tailored to the sample. The current algorithm can be improved based on our findings. For example, these findings indicate that output libraries and candidate oligos are not absolutely exclusive to the target cells used as input in their evolution process. Therefore, the personalized algorithm should include an early step that screens any incoming sample against the library of previously-generated oligonucleotides, to shunt directly to synthesis in case effective and selective candidates are found. The algorithm is still personalized per sample, but such decision trees could tremendously improve its efficiency. We are also currently improving methods for candidate selection from sequencing data, based on parameters orthogonal to abundance.

Although the test case was cancer, our findings highlight the possibility to utilize this rapid platform against other targets, such as antibiotic-resistant bacteria, which are a challenge of increasingly critical importance. Central to this challenge is the profound asymmetry between the period of time required for the development of a new antibiotic and the period of time required for targets to develop resistance to it: while new drugs require on average more than 10 years and $2.6B to develop^34^, antibiotic resistance in bacteria could arise within a few generations, or on the scale of hours^35,36^, rendering the pharmaceutical industry extremely limited in dealing with this challenge. The present platform could offer a significant advantage in this battle. Interestingly, oligonucleotides have the additional unique advantage of being “digitizable” - they can be distributed as electronic sequence files and synthesized locally, owing to their facile synthesis. We have recently shown using network models that this concept formulates the most effective strategy to date to mitigate global pandemics^37^. Coupled with an ultra-rapid discovery system, a tool is created which, arguably, must be pursued.

## Methods summary

Cell lines were purchased from American Type Cell Culture (ATCC). Human primary acute myeloblastic leukemia (AML) cells were isolated from donors by standard procedures (Institutional Review Board Approval numbers 0297-15-TLV & 4573-17-SMC). Human (PDX-derived) triple negative breast cancer cells were a kind gift from B. Dekel, Sheba Medical Center, Israel. Target blasts were isolated by magnetic sorting using a commercial kit (Miltenyi Biotec) according to the manufacturer’s instructions. DNA libraries were designed with a 50-nt random core flanked by 20-nt constant regions and ordered from Integrated DNA Technologies (IDT, 5 umol scale). Randomization was done by hand mixing at IDT. All libraries passed in-house QC of uniformity by HPLC prior to usage. Libraries were reconstituted in ultrapure water at a stock concentration of 1 mM. Sequencing was done on an Illumina NextSeq 500 sequencer using NextSeq 500/550 High Output Kit according to the manufacturer’s instructions. Libraries were clustered on Ion Proton spheres using the Ion OneTouch system, sample preparation instrument, and enriched using Ion OneTouch ES (Enrichment System). Clustered sphere QC was done by Ion Sphere Quality Control Kit according to manufacturer instructions. Sorting was performed on a Becton-Dickinson FACSMelody cell sorter equipped with blue, red, and violet lasers (9 color configuration). Cell imaging was done using fluorescent microscopy with a Chroma-49004 or Chroma-49006 filter cubes and Lumencor Sola SE II 365 illumination. Scans were analyzed using NIS Elements AR_software. Cells were identified by Hoechst staining and apoptosis was determined upon co-location with CellEvent™ Caspase-3/7 Green Detection Reagent labeling. Flow cytometry was performed on a Becton-Dickinson Accuri C6 Plus cytometer equipped with 488 nm and 630 nm lasers, and on a Beckman-Coulter Cytoflex cytometer with a B5-R3-V5 laser configuration. Synthesis of selected oligonucleotides including any modification, for both validation and large scale (>1 mmol) experiments, were done by IDT and LGC Axolabs. All animal procedures were performed in the facilities of Science in Action Ltd. (Rehovot, Israel) (National Ethical Approval number 17-3-113). Animals used in this study were female nude mice 9-10 weeks old. Mice (a total of 8 mice/group) were subcutaneously injected (clipping at approximately 24 h prior to injection) with MDA-MB-231 cells (3×10^6^ cells/mouse) into the right flank, 0.2 mL/mouse (using tuberculin syringe with 27 G needle). E8 was injected S.C, once/2 days at 100 mg/kg dose for a period of 11 days. The administration was performed at a constant volume dosage based on individual body weights using a 1 mL insulin syringe with 30 G needle. *Ex-vivo* organ cultures (EVOC) were prepared by Curesponse Ltd. (IRB approval number 0656-18-TLV), stained with H&E, and analyzed by 2 blinded pathologists. Statistical analysis was performed by student’s *t* test assuming equal variances.

## Acknowledgments

The authors wish to thank Dr. Benjamin Dekel (Sheba Medical Center, Ramat Gan, Israel) for the kind gift of TNBC9 cells; to Dr. Seth Salpeter, Dr. Vered Bar, and Ms. Sarah Baum (Curesponse Ltd, Tel Aviv, Israel) for EVOC experiments; to Dr. Anat Globerson-Levin for technical assistance with *in-vivo* biodistribution experiments; and to the entire team at Augmanity and Aummune Ltd. for valuable technical assistance and discussions.

## Author contribution

The following authors designed experiments, performed experiments, analyzed data, and wrote the manuscript: NM, YA, EL, IB. The following authors designed experiments, performed experiments, and analyzed data: NS, LB, GH, GL. The following authors performed experiments and analyzed data: ED, MG, AZ, AP, NR. IA and AS designed experiments and provided valuable materials. AAH and GN provided valuable technical assistance and oversight support. IB oversaw the project.

## Competing financial interests

The authors declare competing financial interests as follows. All authors are employees and shareholders in companies that develop technologies described in this article (NM, LB, GH, IR, AZ, GL, AAH, IB at Augmanity; EL, NS, ED, AP, YA, GN at Aummune). The following authors are listed as inventors on patent applications related to technologies described in this article: IB, NM, IR, GH, YA, EL, AAH (PCT/IB18/00418, pending); IB, NM, EL, LB, ED, IR, GH, YA, AAH (PCT/IB18/00613, pending); IB, NM, AAH, YA, LB, ED, EL, IR, NS (62/738,235, pending).

